# HiTaxon: A hierarchical ensemble framework for taxonomic classification of short reads

**DOI:** 10.1101/2023.09.11.557224

**Authors:** Bhavish Verma, John Parkinson

## Abstract

Whole microbiome DNA and RNA sequencing (metagenomics and metatranscriptomics) are pivotal to determining functional roles within microbial communities. A key challenge in analysing these complex datasets, typically composed of tens of millions of short reads, is accurately classifying reads to their taxon of origin. Traditional reference-based short-read classification tools are compromised by reference database biases, leading to interest in classifiers leveraging machine learning (ML) algorithms. While ML classifiers have shown promise, they still lag reference-based tools in species-level classification. To address this performance gap, attention has turned to approaches that incorporate the hierarchical structure of taxonomic classifications, albeit with limited results. Here we introduce HiTaxon, a hierarchical framework for creating ensembles of reference-dependent and ML classifiers. HiTaxon facilitates data collection and processing, reference database construction and model training to streamline ensemble creation. We show that databases created by HiTaxon improve the species-level performance of reference-dependent classifiers, while reducing their computational overhead. Additionally, through exploring hierarchical methods for HiTaxon, we highlight that our custom hierarchical ML approach improves species-level classification relative to traditional strategies. Finally, we demonstrate the improved performance of our hierarchical ensemble over current state-of-the-art classifiers in species classification using datasets comprised of either simulated or experimentally-derived reads.

## BACKGROUND

Microbiota in the human gut are increasingly being shown to play a key role in health and disease^1^. To understand how they contribute to disease, attention has turned to interrogating their functional capacity using whole microbiome DNA (‘metagenomics’) and RNA (‘metatranscriptomics’) sequencing^2^. A critical component of these analyses is assigning sequenced reads to their taxon of origin^3,4^. While assembling reads into larger contiguous sequences (‘contigs’) can improve taxonomic classification^5^, there are challenges associated with insufficient coverage^6^ and formation of chimeric contigs^7^. This underscores the need for taxonomic classifiers designed specifically for short reads. Traditional reference-dependent tools used for taxonomic analysis can be categorized into marker-based, DNA-Based, or protein-based approaches^8^. In the former, taxon-specific marker genes are used to generate a taxonomic profile for the target dataset^8^. In the latter two methods, reads are mapped to reference databases of DNA or protein sequences for taxonomic classification^8^, thereby informing on the contributions of individual taxa to microbiome function.

One of the best performing DNA-based tools is Kraken2^9^, a k-mer based approach in which each k-mer from a query read is assigned to the lowest common ancestor (LCA) associated with the set of genomes within its reference database, that match that k-mer. A major challenge for Kraken2 and other reference-based approaches is the choice of database used for comparisons. For example, the LCA evaluation scheme used by Kraken2, which improves precision of predictions, hinders species classification when the tool is paired with large reference databases^10^. This arises due to the presence of a large number of identical sequences in related taxa^10^. This issue will only increase as reference databases continue to grow exponentially^11^ and is further confounded by the overrepresentation of a limited number of taxa. Of the 639,981 high-quality bacterial assemblies in the European Nucleotide Archive in November 2018, over 90% were associated with only 20 unique species^12^. This, in turn, can be a frequent source of false positives^13^. To mitigate the effects of this compositional bias on taxonomic classification, recent work has shown that creating custom reference databases specific to the environment of interest substantially improves the performance of Kraken2^14^. Together these findings establish the need for a framework capable of systematically curating custom databases for reference-dependent tools appropriate for the dataset being analysed.

As an alternative to reference-dependent classifiers, researchers have explored the application of machine learning algorithms such as Naïve Bayes and Support Vector Machines for taxonomic classification^15,16^. Training these algorithms on manually curated sequence-derived k-mer composition profiles have shown promise in small-scale environments, however, they tend to underperform, relative to reference-dependent tools, in large-scale evaluations^16^. More recent applications have focused on deep learning architectures previously applied to computer vision^17^ and natural language processing^18,19^. While deep learning models demonstrate comparable or even superior performance to reference-dependent classifiers at higher ranks of taxonomy, they remain limited in their ability to perform species-level assignments.

Given algorithmic differences between machine learning classifiers and reference-dependent tools, attempts have been made to develop ensemble classifiers^19-22^. Although ensembles were shown to improve performance, evaluations were either conducted at the genus-level^19^ or involved classifiers trained to discriminate between fewer than a few hundred species^20-22^. Further innovations for short-read classification include the use of integrating taxonomic structural hierarchy but these have been restricted to genus-level classification^19,23^. Here we present HiTaxon, a hierarchical ensemble that combines reference-dependent and machine learning classifiers to accurately perform species-level classifications. Through streamlining data collection and processing, database construction, model training, and ensembling, HiTaxon generates custom classifiers appropriate for the dataset of interest (Figure 1). We show that databases constructed by HiTaxon improved species-level assignments, while also dramatically reducing computational overhead. Adopting a unique approach based on hierarchical machine learning architectures, we further demonstrate improved species-classification over existing hierarchical machine learning techniques. Finally, we show that ensembling machine learning classifiers with Kraken2 using a hierarchical classification framework, improved performance over any other classifier.

**Figure 1.**
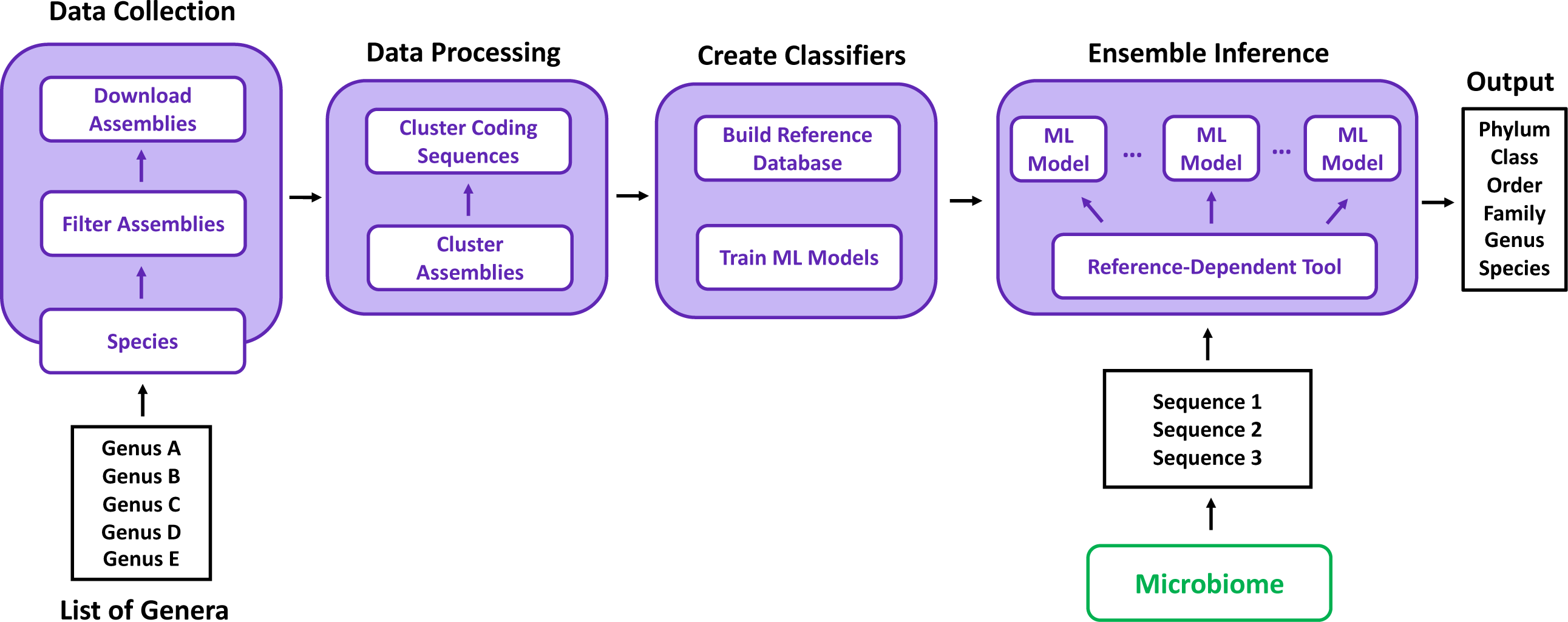
Overview of HiTaxon. Given a list of genera, HiTaxon uses pre-set filters to download high-quality assemblies for species encompassed in the genera. A two-step clustering approach is employed to reduce redundant genomic information, creating species-specific non-redundant sets of coding sequences. These sets of sequences can be used to create a hierarchical ensemble of reference-dependent and machine learning classifiers.

## MATERIALS AND METHODS

### Reference data collection and processing

To create reference databases for reference-dependent classifiers and train ML models, HiTaxon filters RefSeq (accessed August 24th, 2022) for all assemblies encompassed within a query set of genera. On a per-species basis, HiTaxon compiles a list of assemblies, prioritizing higher-level assemblies. At each assembly level, HiTaxon uses the RefSeq category assigned to each assembly as a secondary selection attribute. Using this priority-based ordering, HiTaxon aims to generate species-specific lists composed of 10 high quality assemblies. In instances where a species is represented by fewer than 10 assemblies in RefSeq, all assemblies are added to the list. However, when a species is represented by more than 10 assemblies, assemblies of the lowest quality (i.e worst combination of assembly level and RefSeq category) that represent strains in which better quality assemblies had been obtained are excluded. Coding sequences associated with this list of high-quality assemblies are downloaded using the NCBI Datasets command-line tool^11^ (v. 15.3.1). Post-download, HiTaxon used fastANI^24^(v. 1.33) to compare coding sequences in each assembly to calculate average nucleotide identities (ANI). Assemblies are then grouped based on ANI scores using the OPTICS clustering algorithm^25^ (scikit-learn v. 1.1.3). Adopting this ANI mediated unsupervised clustering approach avoids inconsistencies that can otherwise arise when relying only on a nomenclature based approach^26^. For each group of assemblies generated by OPTICS, the assembly with the highest N50 score, as provided by RefSeq, is selected as the representative assembly. Subsequently, HiTaxon aggregates individual coding sequences into a single species-specific FASTA file. CD-HIT-EST^27^ (v. 4.8.1) is then applied on the aggregated FASTA file to cluster individual coding sequences sharing 99% sequence identity, creating a species-specific dataset of non-redundant coding sequences.

### Independent Classifiers

Given that hierarchical machine learning has shown promise for improving classification performance^19,23^, we investigated four hierarchical frameworks (Figure 2): 1) a local classifier per level (LCL) approach^28^ that uses a single classifier for each taxonomic level; 2) a hierarchy informed LCL approach^28^, in which classifications at higher ranks determine the subset of predictions considered possible for lower ranks; 3) a local classifier per parent node (LCPN) approach^28^ that expands on the hierarchy informed approach through training multi-class classifiers for each taxon across all taxonomic ranks and uses the output of higher-taxonomic ranked parent models to determine which child model is used for subsequent classifications at the lower rank; and 4) a hybrid of the LCL and LCPN approaches in which the LCL method is used for phylum to family level classification, while the LCPN method is used for genus and species level predictions. We note that the LCL approach has previously been used for species level classification^16^. However, prior applications of the LCPN approach to taxonomic classification were either restricted to higher taxonomic ranks^23^, or were reliant on larger genomic fragments as input^29^. To our knowledge, neither the hierarchy informed LCL approach or the hybrid approach have been used for the purposes of taxonomic classification.

**Figure 2.**
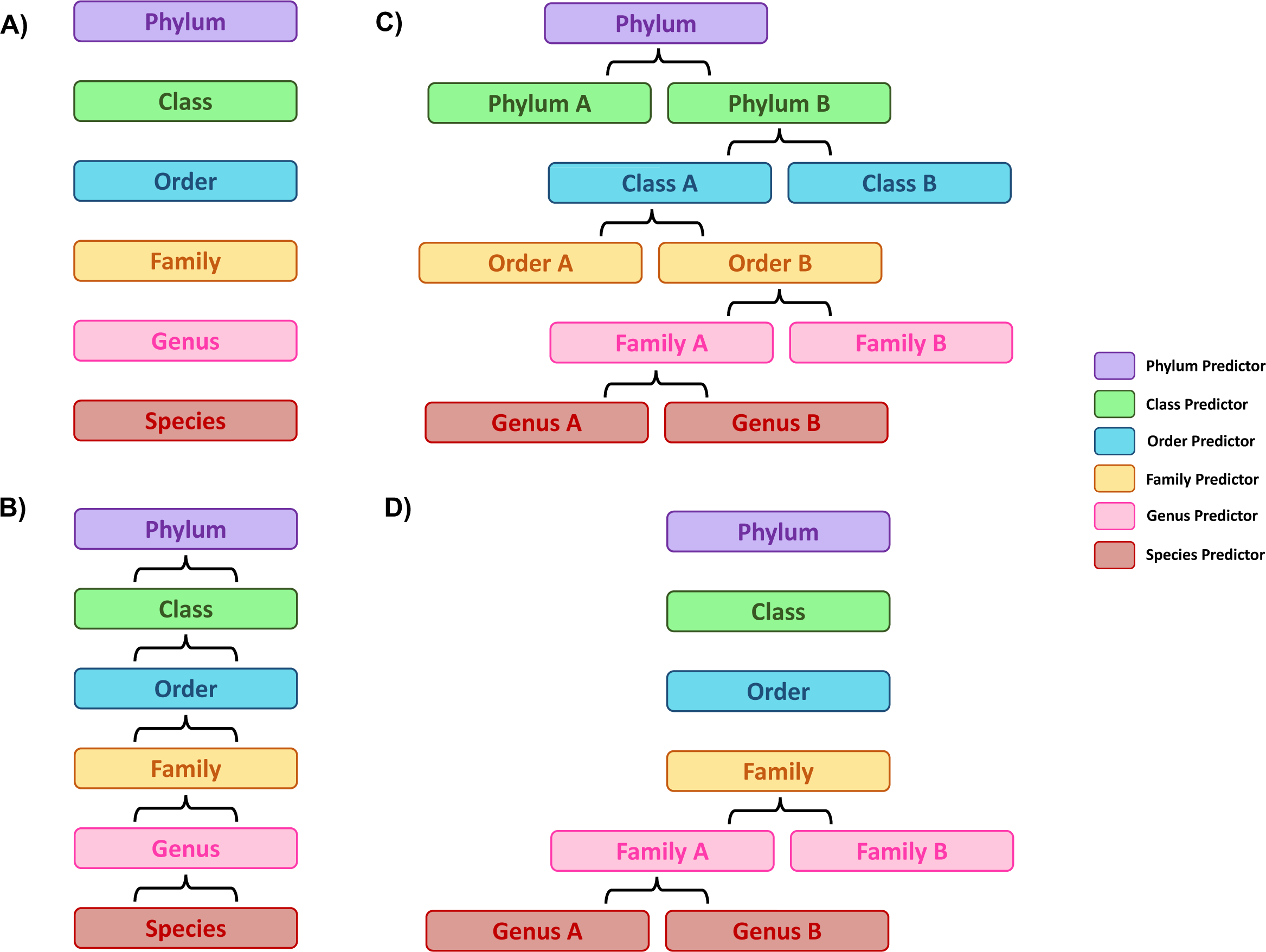
Hierarchical machine learning methods for species classification. (A) The local classifier per level (LCL) approach uses models trained for a specific taxonomic rank to generate predictions at that rank. (B) The hierarchy-informed local classifier per level (hierarchy-informed LCL) method uses the same classifiers employed in the LCL approach, but the outputs derived from higher taxonomic ranks are used to reduce the prediction search space in the subsequent lower taxonomic level. (C) The local classifier per parent node (LCPN) method uses predictions at higher taxonomic ranks to determine which child classifier to use at lower taxonomic levels. (D) The LCL-LCPN method is a custom hybrid approach which uses the LCL strategy from phylum to family and the LCPN protocol from genus to species.

FastText^30^(v. 0.9.2), a CPU-efficient NLP word embedding algorithm, was used as the core supervised ML algorithm for the four hierarchical approaches. In addition to its competitive performance with convolutional and recurrent neural networks, we used FastText because its shorter training time makes it better amenable to the multi-classifier nature of these local hierarchical architectures. Furthermore, a prior application of FastText to taxonomic classification reported competitive performance to older alignment based algorithms^31^. In addition to the ML classifier, we also investigated the performance of two state-of-the-art reference-dependent classifiers, Kraken2^9^ (v. 2.1.2) and Kaiju^32^ (v. 1.9.2) to determine which tool to use as the primary reference-dependent classifier for HiTaxon.

### Ensembles

Ensembles of Kraken2 and Kaiju with ML classifiers used the phylum to genus-level predictions of reference-dependent classifiers. These genus-level outputs were used to determine which ML classifiers generated predictions at the species-level. If an ML prediction had a softmax score less than 0.5, the species-level prediction of the reference-dependent tool was used instead. We also generated an ensemble combining Kaiju with Kraken2, in which Kraken2’s predictions were used from the phylum to species-level, however, in instances where Kraken2 was unable to generate a species-level output, Kaiju’s phylum to species-level prediction was used instead.

### Classifier Evaluation

To evaluate the performance of individual classifiers and ensembles we simulated multiple datasets that correspond to the genus-level identity of different microbiome environments (human gut, human conjunctival, marine surface, rumen), using sequences that were excluded from RefSeq to avoid data leakage (Table 1 and S1 Text). In addition, we also tested the performance of each classifier on gold-standard experimental datasets of DNA and RNA reads in which the taxa responsible for the reads were known (Table 1 and S1 Text).

**Table 1.** Characteristics of the training and test data used for taxonomic classifiers.

| Dataset Name | Training Genera | Training Species | HiTaxon-DB Coding Sequences | RefSeq-DB Coding Sequences | Testing Species | Testing Reads | Simulated |
| --- | --- | --- | --- | --- | --- | --- | --- |
| Human Gut | 58 | 1718 | 12,027,988 | 269,431,618 | 522 | 3,415,471 to 3,605,377 | Yes |
| Human Conjunctival | 10 | 780 | 5,487,293 | NA | 256 | 1,996,771 to 2,110,540 | Yes |
| Marine Surface | 43 | 1,651 | 11,358,799 | 182,996,521 | 337 | 2,867,107 to 3,083,814 | Yes |
| Rumen | NA | NA | NA | NA | NA | 1,159,325 | Yes |
| Mock Community 1 | 8 | 646 | 6,018,839 | 205,425,935 | 8 | 4,556,549 | No |
| Mock Community 2 | 23 | 1275 | 10,863,883 | 389,602,189 | 28 | 2,599,072 | No |

To create custom Kraken2 and Kaiju HiTaxon databases (Kraken2-HiTaxon-DB, Kaiju2-HiTaxon-DB) suitable for a specific dataset, we used species-specific HiTaxon curated coding-sequences encompassed in the genera relevant to that specific dataset (Table 1). Note that we only focused on creating non-redundant coding sequences for NCBI formal named species that adhere to the international code of nomenclature of prokaryotes (ICNP). Additionally, for most datasets, we also created databases for both Kraken2 and Kaiju (Kraken2-RefSeq-DB, Kaiju2-RefSeq-DB) in which a maximum of 5000 assemblies from RefSeq were randomly selected for each species used in the HiTaxon database for that particular dataset to mimic an ad-hoc approach to database construction (Table 1). Since Kaiju is a peptide-based classifier, Prodigal^33^ was used to perform 6-frame translations of the reference nucleotide sequences.

To train the ML classifiers for the hierarchical frameworks, 500,000 150bp paired-end ART^34^ (v. 2.5.8) simulated reads were derived, for each species, from the same set of HiTaxon species-specific coding-sequences used to build HiTaxon-DB databases. Prior to training, these reads were aggregated, merged, tokenized in 13mers, and shuffled. During generation of ML classifiers for the human conjunctival environment, we used 464 RefSeq assemblies that were not selected to be used for training, to create a validation set composed of 1,221,740 reads. This validation set was then used to select optimal hyperparameters for the LCL species-level FastText classifier, through a random search approach^35^. Based on this search, we found that a FastText model with an embedding dimension of 200, a context window of 5, wordNgrams of 1, and a learning rate of 0.5 that was trained for 15 epochs and evaluated using a confidence threshold of 0.5 provided the best species-level performance. The same set of hyperparameters (excluding learning rate and number of epochs such that training is kept under 24 hours) were used for the remaining classifiers and subsequent datasets to best simulate the application of HiTaxon using default settings.

Classifier performance was assessed using the following metrics:

Macro Precision = Mean of *TP/(TP + FP*) across all species

Macro Recall = Mean of *TP/(TP + FN)* across all species

Macro F1 = Mean of *2(Precision x Recall)/(Precision + Recall)* across all species

### Computing Environment

Classifiers were created and benchmarked on the SciNet compute system Niagara, using nodes with two Intel(R) Xeon(R) Gold 6148 CPU @ 2.40GHz with 202 GB RAM running CentOS 7.

## RESULTS

### Overview of HiTaxon

Here we were interested in addressing two major concepts associated with taxonomic classification accuracy. First, given their known capacity to significantly impact classifier performance, the reference database used for taxonomic assignments; and second, the use of a hierarchical classification architecture for an ensemble of reference-dependent and ML classifiers as this could both improve species classification and allow for the assignment of labels to higher ranks for lower confidence predictions to better avoid false positives. To integrate these concepts, we developed the HiTaxon framework (Figure 1). For each species of interest, the HiTaxon framework automatically collates a set of non-redundant coding sequences, which is later used to construct an appropriate custom reference database. Following this, HiTaxon trains multiple local FastText classifiers for species prediction. Finally, HiTaxon employs a hierarchical classification algorithm to integrate the predictions of Kraken2 with the trained ML models. In the following, we assess the performance of the HiTaxon framework, including the custom database generation and hierarchical classifier relative to other classifiers.

### Non-redundant coding sequences elevate reference-dependent species classification

Key to the performance of any classifier is the generation of appropriate reference databases. Given a list of genera, HiTaxon generates sets of non-redundant coding sequences for all encompassed NCBI formal named species.

We benchmarked the performance of HiTaxon’s systematic database curation approach by pairing Kraken2^9^ and Kaiju^32^, with either unprocessed RefSeq sequences (RefSeq-DB) or HiTaxon curated sequences (HiTaxon-DB) and applied these classifiers to taxonomically assign simulated sequence reads. Classifiers using the RefSeq-DB and HiTaxon-DB databases were built with, respectively, 67,490 and 8,407 assemblies (269,431,618 coding sequences and 12,027,988 coding sequences) for 1,718 species encompassed within 58 genera associated with the human gut microbiome^36^. Using 1,641 assemblies excluded from RefSeq that comprise of 522 species encompassed in the same 58 genera, we simulated 10 unique test sets that ranged between 3,415,471 to 3,605,377 paired-end reads.

Focusing on species-level performance (Figure 3A), we found that Kraken2 outperformed Kaiju and the use of HiTaxon-DB outperformed the RefSeq-DB database (Kraken2-HiTaxon-DB: 0.65 Macro F1 ± 0.0039; Kaiju-HiTaxon-DB: 0.54 Macro F1 ± 0.0037; Kraken2-RefSeq-DB: 0.62 Macro F1 ± 0.0042; and Kaiju-RefSeq-DB: 0.51 Macro F1 ± 0.003; Wilcoxon Signed-Rank test with Benjamini-Hochberg correction: p < 0.05). Performance gains were largely driven through fewer unclassified reads; Kraken2 and Kaiju assigned 326,199.6 ± 16,773.9 and 357,027.0 ± 11,461.8 more reads respectively using the HiTaxon-DB database. These results highlight previous findings in which the additional noise introduced with redundant sequences compromises the ability of tools that utilize LCA evaluation algorithms to classify reads at the level of species^10^.

**Figure 3.**
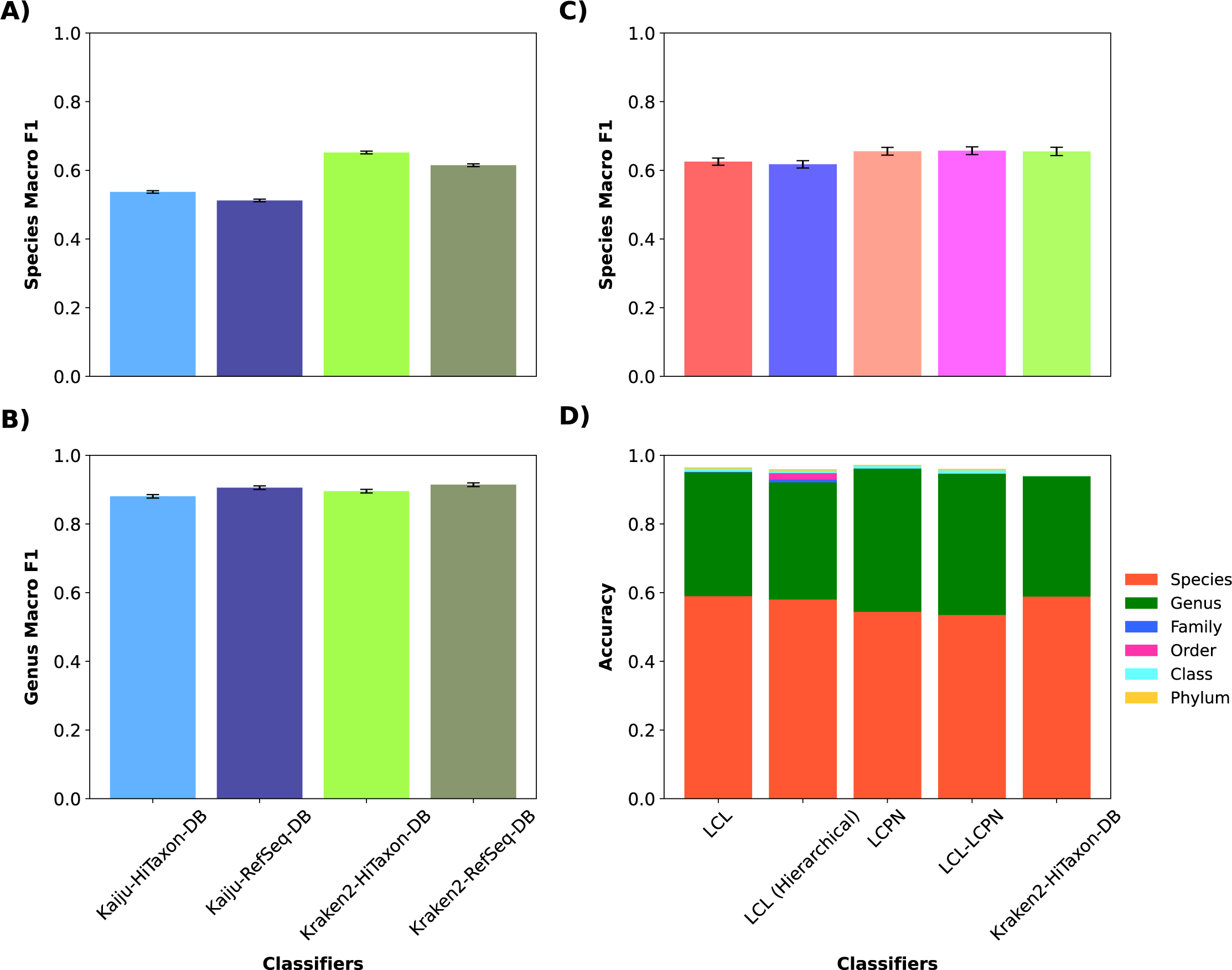
Performance of individual classifiers. Using simulated test sets consistent with the genera-level identity of the human gut microbiome, the (A) species and (B) genus-level macro F1 score was computed for classifiers with the RefSeq-DB database and the HiTaxon-DB databases. Using simulated test sets consistent with genera-level identity of the human conjunctival microbiome, the (C) species macro F1 score and (D) accuracy across all taxonomic ranks was computed for each hierarchical machine learning approach and Kraken2-HiTaxon-DB.

With respect to genus classification (Figure 3B), Kraken2 and Kaiju performed similarly. However, both Kraken2-HiTaxon-DB and Kaiju-HiTaxon-DB (Kraken2: 0.90 Macro F1 ± 0.0052, Kaiju: 0.88 Macro F1 ± 0.0050) exhibited slightly poorer performance relative to the use of the RefSeq-DB database (Kraken2-RefSeq-DB: 0.92 Macro F1 ± 0.0054, Kaiju-RefSeq-DB: 0.91 Macro F1 ± 0.0053; Wilcoxon Signed-Rank test with Benjamini-Hochberg correction: p < 0.05). Unlike species classification, we found that Kraken2-RefSeq-DB and Kaiju-RefSeq-DB left fewer reads unclassified at the genus-level relative to the HiTaxon-DB database (58,733.7 ± 3,154.90 and 20,917.2 ± 15,050.0 additional reads identified using Kraken2-RefSeq-DB and Kaiju-RefSeq-DB respectively). These results again highlight the sensitivity of LCA evaluation schemes to reference databases. This marginal improvement extended to higher taxonomic ranks (S1 Data). The narrowing of performance between Kaiju and Kraken2 likely reflects the ability of synonymous SNPs to better discriminate at the level of species.

Beyond improved species-classification performance, the HiTaxon-DB database approach provides substantial benefits from the compute perspective (Table 2), as it required a reduced amount of memory (HiTaxon-DB: 63GB and 11GB vs. RefSeq-DB: 333GB and 210GB for Kraken2 and Kaiju respectively). Additionally, the time needed to construct the HiTaxon-DB databases was lower than for the RefSeq-DB databases (HiTaxon-DB: 29 minutes, 18 minutes vs. RefSeq-DB: 13 hours and 27 minutes, 6 hours and 31 minutes for Kraken2 and Kaiju respectively).

**Table 2.** Computing resources required for Human Gut databases created for Kaiju and Kraken2.

| **Tool** | **Database** | **Build Time** | **Memory** |
| --- | --- | --- | --- |
| Kaiju | RefSeq-DB | 6 Hours and 31 Minutes | 210 GB |
| Kaiju | HiTaxon-DB | 18 Minutes | 11 GB |
| Kraken2 | RefSeq-DB | 13 Hours and 27 Minutes | 333 GB |
| Kraken2 | HiTaxon-DB | 29 Minutes | 63 GB |

### A hybrid approach to hierarchical machine learning improves species-level performance

Having demonstrated the improved performance of the custom database for species-level classification with existing algorithms, we next investigated how a ML algorithm might benefit from such preprocessing. Since adoption of hierarchical frameworks, reflecting different levels of taxonomy, have shown potential for improving predictive performance, we investigated four hierarchical strategies: LCL, Hierarchy-informed LCL, LCPN, and LCL-LCPN. To compare hierarchical strategies, we trained each approach to classify 780 species belonging to 10 genera associated with the human conjunctival microbiome^37^. Species-specific HiTaxon curated coding sequences used to train these architectures were obtained from 3,648 RefSeq assemblies (5,487,293 coding sequences). Benchmarking of each approach was evaluated using 10 unique test sets, ranging between 1,996,771 to 2,110,540 paired-end reads, derived from 767 RefSeq excluded assemblies and associated with 256 species encompassed in the 10 human conjunctival genera. In addition, we also built a HiTaxon-DB database for Kraken2, the best performing species classifier in the previous test, using the same set of HiTaxon curated coding sequences utilized for training the ML models.

Of the four strategies, the two LCPN-based approaches performed the best in terms of species-level assignments (Figure 3C: LCL-LCPN-0.657 macro F1 ± 0.0112; LCPN - 0.655 macro F1 ± 0.0113 vs LCL - 0.625 macro F1 ± 0.0105; and Hierarchy-informed LCL - 0.617 macro F1 ± 0.01055; Wilcoxon Signed-Rank test with Benjamini-Hochberg correction: p < 0.05). Between the two LCPN-based approaches, the hybrid LCL-LCPN method had slightly better species performance. When comparing the performance of the LCPN and LCL-LCPN methods at the family-level, recalling that the hybrid LCL-LCPN approach uses the LCL classifier for family predictions, we found that the LCL-LCPN approach was 2.23% greater on average (Figure 3D). This reduced performance at the family level by the LCPN method suggests that compounding errors from incorrect predictions at higher ranks outweigh the benefit of using hierarchy-informed predictions at all taxonomic levels. This suggestion is further strengthened when comparing the species-level performance of the two LCL approaches. Despite using the same trained classifiers, the hierarchy-informed LCL method performed worse at the species-level relative to the conventional LCL strategy. When compared to Kraken2, the LCL-LCPN approach yielded a minor yet significant improvement in species-level classification (LCL-LCPN: 0.657 macro F1 ± 0.0112, Kraken2: 0.655 macro F1 ± 0.012; Wilcoxon Signed-Rank test with Benjamini-Hochberg correction: p < 0.05), highlighting that the trained ML classifiers used in our LCL-LCPN method is competitive with current state-of-the-art.

### A hierarchical ensemble with Kraken2 outperforms all other classifiers on simulated reads

Having created a novel hierarchical ML classifier that appears competitive with Kraken2, we reasoned that since the two methods utilize different algorithms, their combination might lead to further performance gains. We therefore generated an ensemble approach adopting a strategy similar to the LCL-LCPN method. Given that Kraken2 outperformed all hierarchical approaches at all ranks above species in the human conjunctival test set, from a macro F1 perspective (S2 Data), we used Kraken2’s genus outputs to determine which species-level ML classifiers to use for predictions. We opted to use ML classifiers as the primary species predictor within our hierarchical ensemble to mitigate classification biases that are associated with over-represented species in reference databases^13^. Despite HiTaxon using a systematic approach to reduce redundant data, we still found that our custom databases for reference-dependent tools resulted in an uneven distribution of sequence data for individual species. For instance, we identified 250,605 coding sequences for *Escherichia coli* and only 4,253 sequences for *Escherichia whittamii*. HiTaxon still provides a significant removal of bias when we consider the original set of sequences, capped at 5000 assemblies per species, comprise 25,332,170 and 8,509 coding sequences for *E. coli* and *E. whittamii* respectively. However, prior to training species-level ML classifiers, we balanced our training set such that an approximately equal number of simulated reads from HiTaxon curated species-specific sequences were used for training; thus ensuring models were not biased towards certain species within particular genera. In instances, where the ML classifier reported low quality predictions (softmax score < 0.5), Kraken2 predictions were used instead.

In addition to evaluating the performance of the ensemble (Kraken2-HiTaxon-Ens) against Kraken2, Kaiju and the LCL-LCPN method, we also investigated two other ensembles involving Kaiju predictions (see Methods). Classifiers were constructed from sequence data pertaining to 1,651 species present in the 43 genera associated with the marine surface microbiome^38^; two reference databases were used, HiTaxon-DB (4,770 assemblies; 11,358,799 coding sequences) and RefSeq-DB (43,807 assemblies; 182,996,521 coding sequences). Unlike, the RefSeq-DB database used for the human gut microbiome, we built the marine surface RefSeq-DB database without capping it to 5000 RefSeq assemblies per species. Despite doing so, the RefSeq-DB marine surface microbiome database used 35.1% fewer RefSeq assemblies when compared to the human gut microbiome database even though it only encompassed 3.6% fewer species (1,651 species vs 1,718); highlighting the reduced coverage of organisms used in the marine surface microbiome simulation. 10 unique test sets composed of 337 species encompassed in the 43 genera used for building reference databases, ranging between 2,867,107 to 3,083,814 paired-end reads, were derived from 974 RefSeq-excluded assemblies and used to compare classifiers (Figure 4A). As before, both Kaiju and Kraken2 exhibited improved performance with the HiTaxon-DB database relative to the RefSeq-DB database – macro F1 scores 0.58 ± 0.0044 vs. 0.53 ± 0.0026 and 0.45 ± 0.0033 vs 0.414 ± 0.003 for Kraken2 and Kaiju respectively (p < 0.05, Wilcoxon Signed-Rank test with Benjamini-Hochberg correction: p < 0.05), illustrating the robustness of HiTaxon’s data reduction framework (S3 Data). Unlike the comparisons involving human conjunctival microbiome sequences, Kraken2-HiTaxon-DB also outperformed the LCL-LCPN approach (macro F1 0.561 ± 0051, p <0.05, Wilcoxon Signed-Rank test with Benjamini-Hochberg correction), suggesting that the LCL-LCPN approach may be compromised by complex datasets.

**Figure 4.**
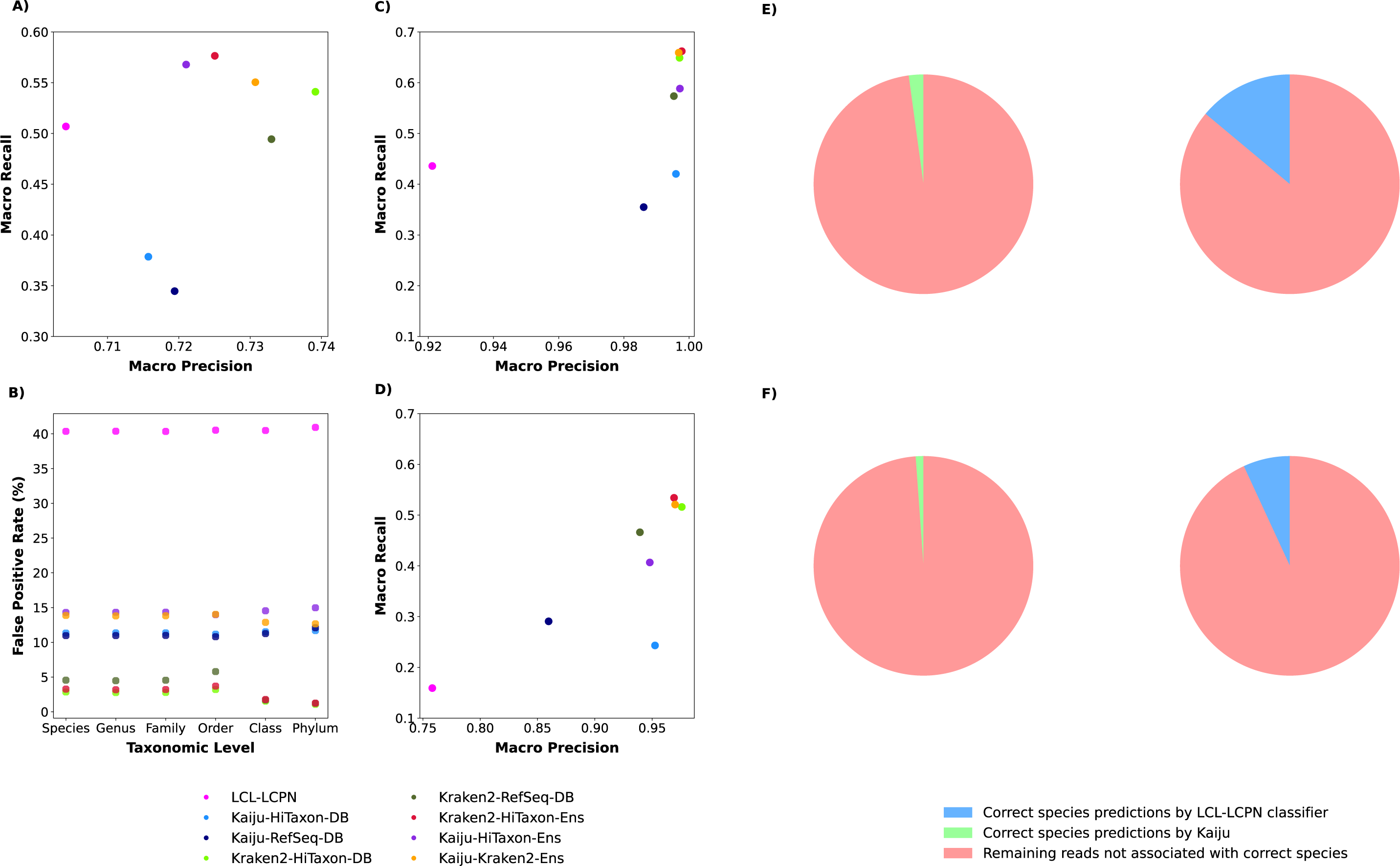
Performance of ensemble classifiers. (A) Using simulated test sets consistent with genera-level identity of the marine surface microbiome, we computed the macro Recall and macro Precision for each classifier. (B) At each taxonomic rank, the proportion of simulated rumen microbiome reads pertaining to taxa not used to create reference databases or train marine surface classifiers that were inappropriately assigned a species label was measured. For both Mock Community 1 (C) and Mock Community 2 (D), which are composed of experimentally-derived reads, species-level macro Precision and Recall were computed. The amount of reads that were not assigned to their correct species by Kraken2 but were correctly predicted by Kaiju-HiTaxon-DB or the LCL-LCPN ML classifier was measured for both Mock Community 1 (E) and Mock Community 2 (F).

Focusing on the ensemble classifiers, we found all three exhibited superior performance to any single classifier, with Kraken2-HiTaxon-Ens performing the best – macro F1 scores 0.609 ± 0.0049, 0.604 ± 0047 and 0.586 ± 0042 for Kraken2-HiTaxon-Ens, Kaiju-HiTaxon-Ens and Kaiju-Kraken2-Ens respectively (p<0.05, Wilcoxon Signed-Rank test with Benjamini-Hochberg correction; S3 Data). Kraken2-HiTaxon also exhibited the highest recall (0.576 ± 0.0053 vs 0.344-0.568 for other classifiers; p<0.05, Wilcoxon Signed-Rank test with Benjamini-Hochberg correction). Given that the primary difference between Kraken2-HiTaxon-Ens and Kraken2-HiTaxon-DB is Kraken2-HiTaxon-Ens’ use of ML classifiers at the species-level, it suggests that their added algorithmic differences and reduced biases allows the hierarchical ensemble to better distinguish between a broader range of species.

### Kraken2 classifiers minimize false positives on unseen taxa

Previous estimates suggest only 2.1% of prokaryotic taxa have been sequenced^39^, suggesting classifiers may be confounded by species not present in their reference datasets and thus resulting in false positives. Although Kraken2-HiTaxon-Ens exhibited the best recall, Kraken2-HiTaxon-DB had higher precision (0.7250 ± 0.0047 vs. 0.739 ± 0.0055; p<0.05, Wilcoxon Signed-Rank test with Benjamini-Hochberg correction: p < 0.05). This motivated us to investigate the prediction of false positives by each classifier, using reads derived from taxa not present within the marine surface microbiome reference database.

From a collection of 349 rumen cultivated genomes^40^ and rumen microbiome metagenome assembled genomes (MAGs)^41^, we simulated a rumen microbiome test set composed of 1,159,325 paired-end reads, comprising 113 species, 57 genera, 33 families, 22 orders, 11 classes, and 6 phyla not present in the marine surface microbiome simulation. At each taxonomic rank, after isolating for those reads in the rumen microbiome test set which were derived from taxa not present in the marine surface microbiome simulation, we measured the amount of species-level false positives that were generated. Overall, we found the proportion of false positives generated by classifiers was relatively consistent across all taxonomic levels (Figure 4B). Relative to other classifiers, the LCL-LCPN approach was most prone to generating false positives (40.5 % ± 0.19 for all taxonomic ranks). Kraken2-HiTaxon-DB performed the best (average of 2.37% ± 0.774 for all taxonomic ranks), indicating that the poor performance of the LCL-LCPN algorithm is not a consequence of HiTaxon’s data collection and processing. We found that Kraken2-HiTaxon-Ens (average of 3.715% ± 0.893 for all taxonomic ranks) was more effective at avoiding false positives relative to Kaiju-Kraken2-Ens (average of 13.5 % ± 0.532 for all taxonomic ranks), second only to Kraken2-HiTaxon-DB (S4 Data).

### Kraken2-HiTaxon improves species classification when applied on experimental reads

To ensure the improved performance of our ensemble classifier translates from simulated datasets to experimentally derived reads, we investigated its ability to classify reads from two experimental studies. The first dataset (‘Mock Community 1’), comprised 4,556,549 paired-end reads, was generated from a sample composed of the eight bacterial taxa associated with the ZymoBIOMICS Microbial Community Standard^42^. Classifiers were created using 6,018,839 HiTaxon curated coding sequences generated from 5,463 assemblies (646 species) that pertain to the eight genera associated with the ZymoBIOMICS taxa. We also created a RefSeq-DB database comprised of 35,809 assemblies (205,425,935 coding sequences) associated with the same 646 species. While Kraken2-HiTaxon-DB performed best among the individual classifiers (macro F1: 0.727), we found its performance improved in combination with both Kaiju and HiTaxon trained ML classifiers (Table 3 and Figure 4C). Further, Kraken2-HiTaxon-Ens performed the best overall (macro F1: 0.759 & 0.738 for Kraken2-HiTaxon-Ens and Kaiju-Kraken2-Ens respectively). Impressively, Kraken2-HiTaxon-Ens exhibited near prefect precision (macro Precision: 0.998). The second dataset (‘Mock Community 2’) comprised of 2,599,072 paired-end and single-end reads from a transcriptomic analysis of human bacterial pathogens^43^. Classifiers were generated using 10,863,883 HiTaxon curated sequences from 9,369 assemblies (1,275 species) that pertain to 23 genera associated with the 28 species in Mock Community 2. As previous, we also created a RefSeq-DB database composed of 100,681 RefSeq assemblies (389,602,189 coding sequences) that are associated with the same 1275 species. Alongside Kraken2-HiTaxon-DB performing best among the individual classifiers (macro F1: 0.622), it’s combination with ML classifiers again provided the best performance (macro F1: 0.651 & 0.626 for Kraken2-HiTaxon-Ens and Kaiju-Kraken2-Ens respectively; Table 3 and Figure 4D). Given both individual Kaiju classifiers outperformed the LCL-LCPN method in isolation (macro F1: 0.243, 0.399 and 0.352 for LCL-LCPN, Kaiju-RefSeq-DB and Kaiju-HiTaxon-DB respectively), we were interested in further investigating the improved performance of Kraken2-HiTaxon-Ens over Kraken2-Kaiju-Ens.

**Table 3.**
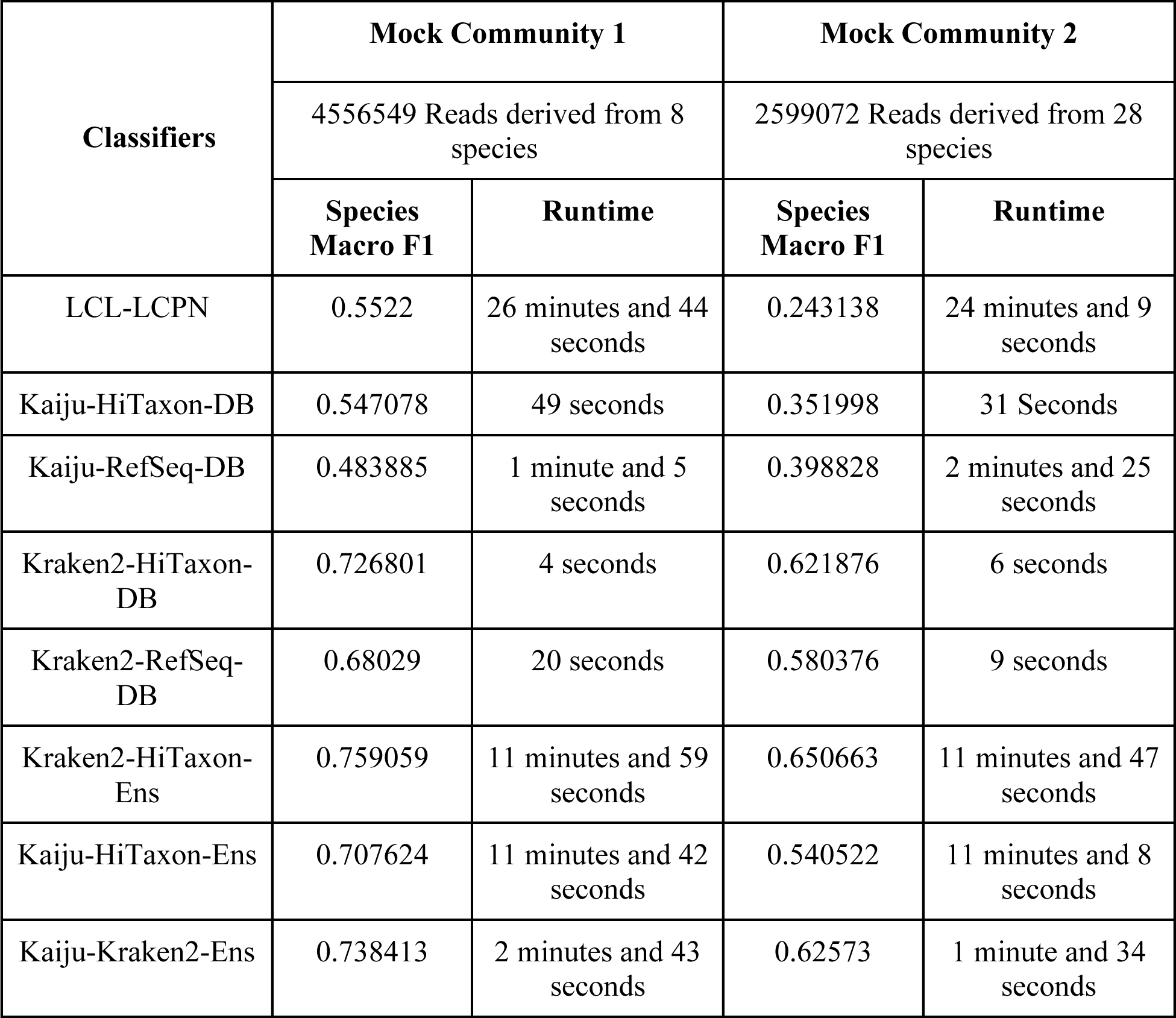
Runtime of classifiers across Mock Community 1 & 2.

A crucial principle for creating a successful ensemble is to incorporate classifiers that offer varied predictions^44^. Consequently, to determine whether Kraken2-HiTaxon-Ens’ superior performance relative to Kaiju-Kraken2-Ens could be attributed to this principle, we measured the performance of Kaiju-HiTaxon-DB and the LCL-LCPN approach on reads from both mock communities that Kraken2-HiTaxon-DB had failed to annotate to the correct species. For Mock Community 1, the LCL-LCPN approach correctly classified more reads than Kaiju-HiTaxon-DB (LCL-LCPN: 197,507 reads; Kaiju-HiTaxon-DB: 30,575 reads) (Figure 4E) and outperformed Kaiju-HiTaxon-DB across all 8 species (S5 Data). Although this might be expected given the improved performance of the LCL-LCPN method observed for Mock Community 1 (Table 3), this performance on reads misassigned or unassigned by Kraken2-HiTaxon-DB also translated to Mock Community 2 (LCL-LCPN: 81,088 reads; Kaiju-HiTaxon-DB: 13,430 reads) (Figure 4F). Further, the LCL-LCPN method classified more reads correctly for 20 of the 28 species associated with this dataset (S6 Data).

These results highlight that, while Kaiju is a better individual predictor in certain instances relative to the LCL-LCPN approach, its predictions exhibit greater overlap with those of Kraken2 and consequently contribute less additional information relative to the ML classifiers used in the Kraken2-HiTaxon-Ens ensemble.

## DISCUSSION

Metagenomics and metatranscriptomics represent powerful technologies to functionally interrogate microbiomes^2^. A key challenge is classifying sequenced reads to their taxa of origin. Here, we present HiTaxon, a framework for generating hierarchical ensembles of reference-dependent and ML classifiers to improve species prediction. A core benefit of HiTaxon is the automated generation of non-redundant coding sequences for database construction and model training. Consistent with previous studies, we found with few exceptions, that the use of these custom databases enhanced species-level predictions^10,14^. Additionally, HiTaxon databases generated fewer false positives relative to databases composed of unprocessed RefSeq sequences. Thus, HiTaxon more effectively mitigates previously reported concerns pertaining to the susceptibility of reference-dependent classifiers with custom databases to generating false positives^45^. Furthermore, with the current RefSeq database occupying over 1 TB of storage and requiring several weeks to build for reference-dependent tools^46^, HiTaxon significantly reduces computational overhead through reducing redundant data.

Given that taxonomic categorization involves organizing organisms into distinct hierarchical groups based on evolutionary relationships^47^, prior to ensembling, we investigated the use of hierarchical ML methods for species classification. We determined that our LCL-LCPN strategy provided improved species-level performance relative to other hierarchical ML approaches. Key to settling on the structure of our hierarchical ensemble, was the finding that errors compounded from misassignments at higher taxonomic ranks outweighed the benefit of using hierarchy-informed predictions across all taxonomic ranks. Rather, limiting hierarchy-informed predictions to lower taxonomic ranks yielded superior performance. While Kraken2 continues to lead the field in taxon assignments, we found its performance could be enhanced through a hierarchical ensembling strategy where Kraken2’s genus-level predictions were supplemented with ML classifiers operating at the species-level. Testing of Kraken2-HiTaxon-Ens using simulated and experimental datasets, demonstrated the best species-level assignments in all instances. Relative to the Kaiju-Kraken2 ensemble, we found Kraken2-HiTaxon-Ens’ improved performance arises from the complementary approaches underlying the two classifiers, fulfilling a core tenant to generating an effective ensemble.

Although HiTaxon offers considerable advantages, we do note several caveats. First, in the analysis of the mock communities, the runtime associated with Kraken2-HiTaxon-Ens classifications is considerably higher than for Kaiju or Kraken2 (Mock Community 1: HiTaxon-Kraken2-Ens: 11 minutes and 59 seconds, HiTaxon-Kraken2-DB: 4 seconds, HiTaxon-Kaiju-DB: 49 seconds; Mock Community 2: HiTaxon-Kraken2-Ens: 11 minutes and 47 seconds, Kraken2-HiTaxon-DB: 6 seconds, Kaiju-HiTaxon-DB: 31 seconds; Table 3). In addition, relative to the other classifiers, the use of HiTaxon’s non-redundant sequences resulted in a marginal decrease in performance at higher taxonomic ranks.

Future design improvements should therefore focus on HiTaxon’s data collection and processing pipeline, to improve performance at higher ranks whilst avoiding compromising the accuracy of predictions at the species-level. Further, since RefSeq is historically biased in taxa associated with medical, industrial or agricultural applications, alternative resources such as collections of MAGs hosted by, for example, the MGnify resource^48^, offer potential for generating more appropriate environment-specific classifiers. With respect to the ML component of our hierarchical ensemble, additional exploration of hyperparameters could further enhance performance, as hyperparameters derived for the human conjunctival microbiome simulation were used throughout this study. Beyond hyperparameters, alternative ML algorithms should continue to be explored. For example, with demonstrated benefits in other domains within biology, fine-tuning large language models, pre-trained on DNA or protein sequence data, offer new avenues for taxon classification^49,50^.

## DECLARATIONS

### Ethics approval and consent to participate

Not applicable

### Consent for publication

Not applicable.

### Availability of data and materials

The software is freely available under the GNU public license V3 and can be accessed through https://github.com/ParkinsonLab/HiTaxon

Additionally, test files used to compare classifiers in the study can be found at https://doi.org/10.5281/zenodo.8335901

### Authors contributions

BV performed experiments, BV undertook analyses, BV and JP conceived the project, BV and JP wrote the draft.

## SUPPLEMENTAL INFORMATION

**S1 Data. Human gut microbiome simulation.** Provides ssemblies used for creating and evaluating classifiers with HiTaxon-DB and RefSeq-DB databases, alongside Macro F1 scores for all taxonomic ranks.

**S2 Data. Human Conjunctival Microbiome Simulation.** Provides assemblies used for training, validating, and testing hierarchical machine learning approaches and Kraken2-HiTaxon-DB, alongside Macro F1 scores for all taxonomic ranks.

**S3 Data. Marine Surface Microbiome Simulation**. Provides assemblies used for HiTaxon sequences, RefSeq-DB, and those used to evaluate classifiers, alongside species Macro Precision, Macro Recall, and Macro F1-score.

**S4 Data. Rumen Microbiome Simulation.** Provides datasets used for creating the simulated test set, alongside classifier performance.

**S5 Data. Mock Community 1.** Provides assemblies used for creating HiTaxon sequences, RefSeq-DB, and those used to evaluate classifiers, alongside species Macro Precision, Macro Recall, Macro F1-score. Additionally, the per-species performance of Kaiju-HiTaxon and the LCL-LCPN classifier on reads that were not associated to their correct species by Kraken2-HiTaxon-DB is given.

**S6 Data. Mock Community 2.** Provides assemblies used for creating HiTaxon sequences, RefSeq-DB, and those used to evaluate classifiers, alongside species Macro Precision, Macro Recall, Macro F1-score. Additionally, the per-species performance of Kaiju-HiTaxon and the LCL-LCPN classifier on reads that were not associated to their correct species by Kraken2-HiTaxon-DB is given.

**S1 Text.** Provides additional information pertaining to the generation of test sets and the inference scheme of hierarchical machine learning architectures.

