## Supplementary material for "HiTaxon: A hierarchical ensemble framework for taxonomic classification of short reads": S1 Text

### SUPPLEMENTARY TEXT

#### Generation of simulated test reads for human gut, human conjunctival, and marine surface microbiome

To evaluate classifiers, we used assemblies excluded from RefSeq as the source for test datasets. These assemblies, which were identified by using a pre-filtered GenBank assembly summary (May 28th, 2023), were unseen throughout HiTaxon's collection and processing of RefSeq sequences. Note: assemblies labeled as suppressed in RefSeq were also used as these assemblies were absent from the RefSeq assembly summary. For each simulated environment, a random subset of 10 assemblies were downloaded for each species encompassed in the genera pertaining to that simulated environment using the NCBI Datasets command-line tool (v. 15.3.1)[1]. 10 unique test datasets were created for each simulated environment. To generate a single test dataset, a single assembly for each species was randomly selected to be used in that specific test dataset. A log-normal distribution with a mean of 0 and standard deviation of 1.5[2] was used to randomly assign a fixed number of reads to each species, relative to a total of 1,000,000 reads. For each species, we computed a baseline number of reads for each gene in the assembly, through dividing a fixed value of total reads (which as indicated prior is species-specific) by the number of genomic coding sequences present in the species assembly. Subsequently, for each CDS associated with each species, 150bp paired-end reads were generated using the RNA-Seq simulator, Polyester[3](v. 1.36), with the 'fold change' parameter randomly assigned a value between 0.125 and 8. Paired-end reads derived from all species were merged using fastq-join[4] (v. 1.3.1) and combined to create a single test dataset.

#### Generation of simulated tests reads for rumen microbiome

In order to generate the rumen microbiome test set, we used Hungate genomes[5] (BioProject Acession: PRJNA471733) and Rumen MAGs[6] (BioProject ID: PRJEB31266). For each of the 116 formal-named species, we aggregated their corresponding genomes and MAGs into a single FASTA file. Since information pertaining to coding sequences were not provided (required for the Polyester pipeline), we used ART[7] (v. 2.5.8) to simulate 10,000 150bp paired-end reads, with a mean fragment size of 200 bp and a standard deviation of 10 bp, from each species FASTA file. Paired-end reads derived from all species were merged using fastq-join[4] (v. 1.3.1) and concatenated into a single test dataset.

### Generation of mock community 1 and 2 test sets

Mock Community 1 used 4,556,549 Illumina paired-end reads from eight bacterial species present within the ZymoBIOMICS Microbial Community Standards[8]. To acquire these reads, we accessed data pertaining to the 8 bacterial isolates from the SRA (BioProject ID: PRJEB29504) and employed fastp[9] (v. 0.23.3) for quality filtration, adapter trimming, and read merging. Following this, rRNA was removed using SortMeRNA[10] (v. 4.3.6) with the smr_v4.3 default database

Mock Community 2 used 2,599,072 Illumina reads from a large-scale transcriptomic analysis of human bacterial pathogens[11]. To acquire these reads, we accessed data pertaining to 28 bacterial species from the SRA (BioProject ID: PRJNA638918). However, due to inconsistencies between species with respect to the reads being single-end or paired-end, the parameters of fastp varied on a per-species basis; resulting in a dataset skewed to a small subset of the 28 species. To create a dataset which allowed us to better evaluate classifier performance over the entire range species, we balanced the dataset by randomly selecting 92,824 reads from each species after rRNA filtration, as this corresponded to the maximum number of reads available to the least abundant species.

### Inference schemes for hierarchical machine learning classifiers

For the LCL method, we trained multiple classifiers to discriminate between taxa from phylum to species. Predictions generated by the LCL approach were evaluated bottom-up, terminating at the lowest-level prediction which had a softmax score greater than or equal to 0.5. Remaining upstream taxonomic predictions were altered such that it remained consistent with the lineage of the lowest level prediction. The same classifiers were reused in the hierarchy-informed LCL strategy, however the evaluation scheme was altered. Unlike the LCL method, this approach employed a two-step evaluation scheme. The first step to evaluation in the hierarchy-informed LCL method was to allow upstream outputs to influence downstream predictions. Briefly, this allowed the initial phylum-level output to restrict class predictions to only those that were present in the predicted phylum. The softmax scores pertaining to this subset of class predictions were renormalized and the output with the highest score was used as the class prediction. This was repeated all the way to the species-level. The second step of the evaluation scheme was to explore predictions bottom-up. The lowest-level prediction with a renormalized softmax score greater than or equal 0.5 is used as the final prediction.

The LCPN approach relies on training an initial classifier that could discriminate between all phyla. This was equivalent to the phylum component of the LCL architecture. However, unlike the LCL approach, classifiers at all other levels were trained to discriminate between a subset of members at a particular rank (i.e. an *Escherichia* classifier was trained to only classify species in the *Escherichia* genera). During this process we often encountered situations where higher-level taxa led to only a single taxon at a lower rank (i.e. a genus with only a single species representative). To enable the application of the LCPN method in environments containing novel organisms, we developed binary classifiers for these higher-rank taxa. These classifiers were trained to distinguish between the correct downstream taxa and a negative class which, if possible, corresponded to a random organism belonging to at least the same phylum. During the prediction phase for reads, the LCPN architecture ensured that the reads were exclusively fed as input to classifiers specified by upstream predictions. These are then evaluated in a bottom-up fashion, terminating at the prediction which had a softmax score greater than or equal to 0.5.

The LCL-LCPN method uses the LCL approach from the phylum to family level and the LCPN strategy from the genus to species level. During evaluation, predictions are considered in a bottom-up fashion, terminating at the lowest-level prediction which had a softmax score greater than or equal to 0.5. Remaining upstream taxonomic predictions were altered such that it remained consistent with the lineage of the lowest level prediction.
